# Dual nature of human ACE2 glycosylation in binding to SARS-CoV-2 spike

**DOI:** 10.1101/2020.07.09.193680

**Authors:** Ahmad Reza Mehdipour, Gerhard Hummer

## Abstract

Binding of the spike protein of SARS-CoV-2 to the human angiotensin converting enzyme 2 (ACE2) receptor triggers translocation of the virus into cells. Both the ACE2 receptor and the spike protein are heavily glycosylated, including at sites near their binding interface. We built fully glycosylated models of the ACE2 receptor bound to the receptor binding domain (RBD) of the SARS-CoV-2 spike protein. Using atomistic molecular dynamics (MD) simulations, we found that the glycosylation of the human ACE2 receptor contributes substantially to the binding of the virus. Interestingly, the glycans at two glycosylation sites, N90 and N322, have opposite effects on spike protein binding. The glycan at the N90 site partly covers the binding interface of the spike RBD. Therefore, this glycan can interfere with the binding of the spike protein and protect against docking of the virus to the cell. By contrast, the glycan at the N322 site interacts tightly with the RBD of the ACE2-bound spike protein and strengthens the complex. Remarkably, the N322 glycan binds into a conserved region of the spike protein identified previously as a cryptic epitope for a neutralizing antibody. By mapping the glycan binding sites, our MD simulations aid in the targeted development of neutralizing antibodies and SARS-CoV-2 fusion inhibitors.

## Introduction

Angiotensin converting enzyme 2 (ACE2) is an enzyme catalyzing the hydrolysis of angiotensin II into angiotensin (1–7), which lowers blood pressure (1). A single transmembrane helix anchors this enzyme into the plasma membrane of cells in the lungs, arteries, heart, kidney, and intestines (2). The vasodilatory effect of ACE2 has made it a promising target for drugs treating cardiovascular diseases (3).

ACE2 also serves as the entry point for some coronaviruses into cells, including SARS-CoV and SARS-CoV-2 (4-6). The binding of the spike protein of SARS-CoV and SARS-CoV-2 to the peptidase domain of ACE2 triggers endocytosis and translocation of both the virus and the ACE2 receptor into endosomes within cells (4). The human transmembrane serine protease 2, TMPRSS2, primes the spike protein for efficient cell entry by cleavage at the boundary between the S1 and S2 subunits or within the S2 subunit (4).

The structure of the ACE2 receptor in complex with the SARS-CoV-2 spike receptor binding domain (RBD) (7-9) has identified the major RBD interaction regions as helix H1 (Q24-Q42), a loop in a beta sheet (K353-R357), and the end of helix H2 (L79-Y83). With a 4-Å cut-off, 20 residues from ACE2 are interacting with 17 residues from the RBD, forming a buried interface of ∼1,700 Å^2^ (7).

The structure of full-length ACE2 has been resolved in the presence of a neutral amino acid transporter (B^0^AT1, also known as SLC6A19) (9). ACE2 functions as chaperone for B^0^AT1 and is responsible for its trafficking to the plasma membrane of intestine epithelial cells (10). Although it was speculated that B^0^AT1 prevents ACE2 cleavage by TMPRSS2 and thus could suppress SARS-CoV-2 infection (9, 11), other studies showed that SARS-CoV-2 can infect human small intestinal enterocytes where ACE2 should presumably be in complex with B^0^AT1 (12).

Both the ACE2 receptor and the spike protein are heavily glycosylated. Several glycosylation sites are near the binding interface (7, 9, 13, 14). Therefore, one may anticipate that these glycosylation sites are involved in the binding process. However, so far most of the studies focus on the effect of interactions at the binding interface of ACE2 and the spike protein (15, 16) and the effect of glycosylation on the binding process has been largely overlooked. The only well-characterized position on ACE2 is N90. It is known from earlier SARS-CoV studies that glycosylation at the N90 position might interfere with virus binding and infectivity (17). Also, recent genetic and biochemical studies showed that mutations of N90, which remove the glycosylation site directly, or of T92, which remove the glycosylation site indirectly by eliminating the glycosylation pattern, increase the susceptibility to SARS-CoV-2 infection (18, 19). However, a detailed molecular mechanism by which glycosylation at N90 or other glycosylation sites impacts the host-virus interactions is urgently needed.

We examine the impact of ACE2 glycosylation on SARS-CoV-2 spike binding by performing extensive molecular dynamics (MD) simulations. We find that glycosylation at sites N90 and N322 significantly affects binding of ACE2 to the spike protein. Remarkably, glycans at these sites have opposite effects, interfering with spike binding in one case, and strengthening binding in the other. Our findings provide direct guidance for the design of targeted antibodies and therapeutic inhibitors of viral entry.

## Results

### The RBD is stably bound to ACE2

To obtain molecular insight into the role of ACE2 glycosylation in spike-RBD binding, we performed MD simulations of a homodimeric membrane-anchored ACE2 receptor complexed with two spike RBDs and two B^0^AT1 (Fig. 1*A*). In addition, we simulated several variants of this complex, with and without ACE2 receptor, RBDs, and B^0^AT1 transporter (SI Appendix, Table S1). ACE2 has seven N-glycosylation sites (SI Appendix, Fig. S1) and based on mass-spectrometry (MS) data (13), one O-glycosylation site is also glycosylated. B^0^AT1 carries five glycosylation sites, and the RBD one. For these glycosylation sites, we considered distinct glycosylation patterns: two variants of homogeneous N-glycosylation and three variants of heterogeneous glycosylation (Fig. 1*B* and SI Appendix, Table S1). For each of the 11 setups, we performed 1-µs long MD simulations. In the simulations with RBDs bound to ACE2, their interaction interface remained stable throughout the simulations, as judged by the native contact profiles relative to the electron cryo-microscopy (cryoEM) structure (PDB ID 6M17) (SI Appendix, Fig. S2). The peptidase domain (residues 21-610) of the ACE2 receptor dimer and the RBDs are also internally stable, as judged by the root-mean-square distance (RMSD) relative to the cryoEM structure (PDB ID 6M17) (SI Appendix, Fig. S2). There are two dimerization interfaces between the peptidase domains of the ACE2 monomers in the complex. While the lower interface, containing an extensive network of interactions, remains stable during the simulations, the upper interface, which has only few interactions, is floppy during the simulations (SI Appendix, Fig. S3). Overall, the presence or absence of B^0^AT1 or the different patterns of glycosylation do not have pronounced effects on the conformations of ACE2, the RBD, and their contact interface.

**Figure 1.**
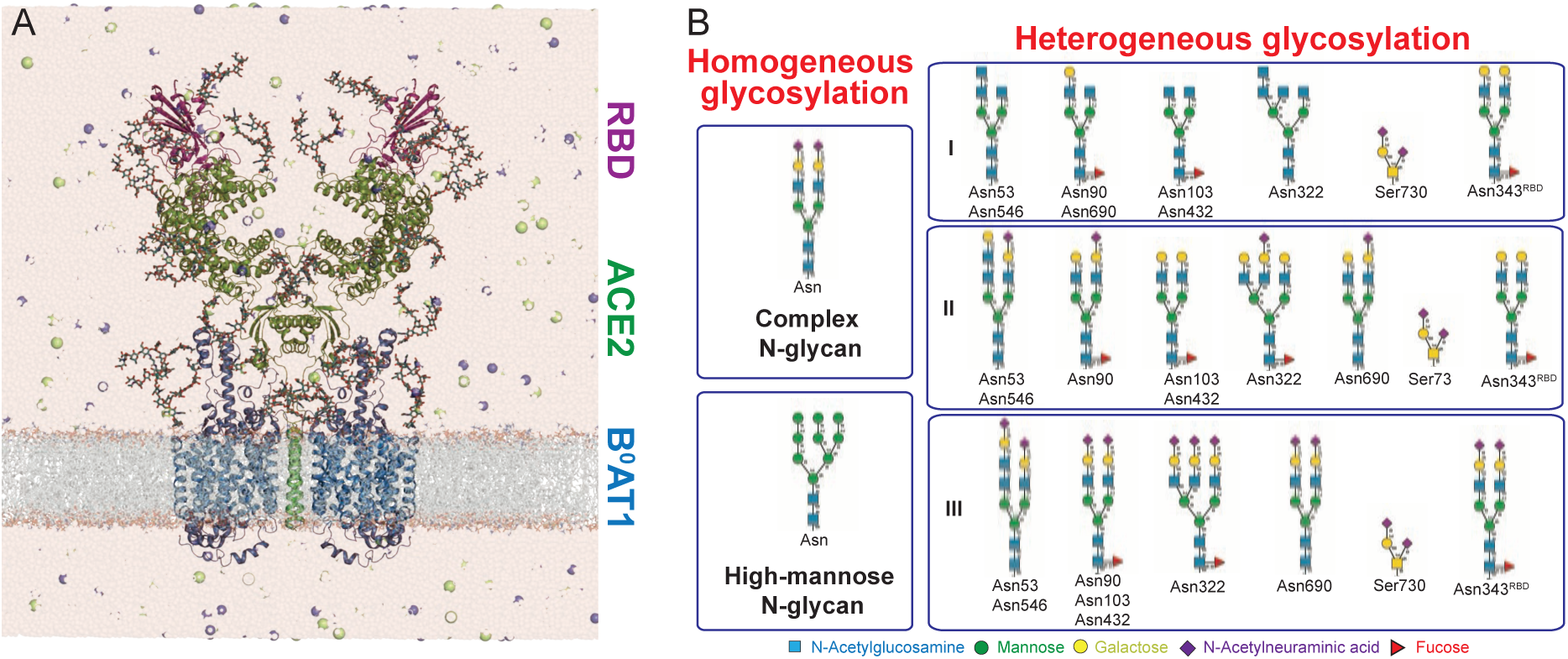
MD simulations of fully glycosylated B^0^AT1-ACE2-RBD complex. (*A*) System setup of the complex with explicit water and physiological concentration of ions. B^0^AT1, ACE2, and the RBD are shown in blue, green, and magenta cartoon, and glycans as licorice. (*B*) Distinct glycosylation patterns used in the simulations: two variants of homogeneous N-glycosylation (left panel) and three variants of heterogeneous glycosylation (right panels).

### The N322 glycan strengthens RBD binding

Geometrically, the glycans at four glycosylation sites (N53, N90, N103, and N322) have the possibility to interact with the RBD. To quantify the interactions of each glycan with the RBD, we calculated the number of residue-residue contacts between each glycan and the RBD in the six simulations where the RBD was present. Note that in each of these simulations we have two copies of the RBD-ACE2 complex, giving us 6×2=12 copies to analyze.

The two glycans at positions N90 and N322 interact most strongly with the RBD (Fig. 2*A* and S3). The N322 glycan has on average between 5 and 7 interactions with the RBD in most of the simulations (in 8 out of the 12 independent RBD-ACE2 complexes) (SI Appendix, Fig. S4). In the remaining four complexes, inter-glycan interactions with the N546 glycan drive the N322 glycan away from the RBD (SI Appendix, Fig. S5).

**Figure 2.**
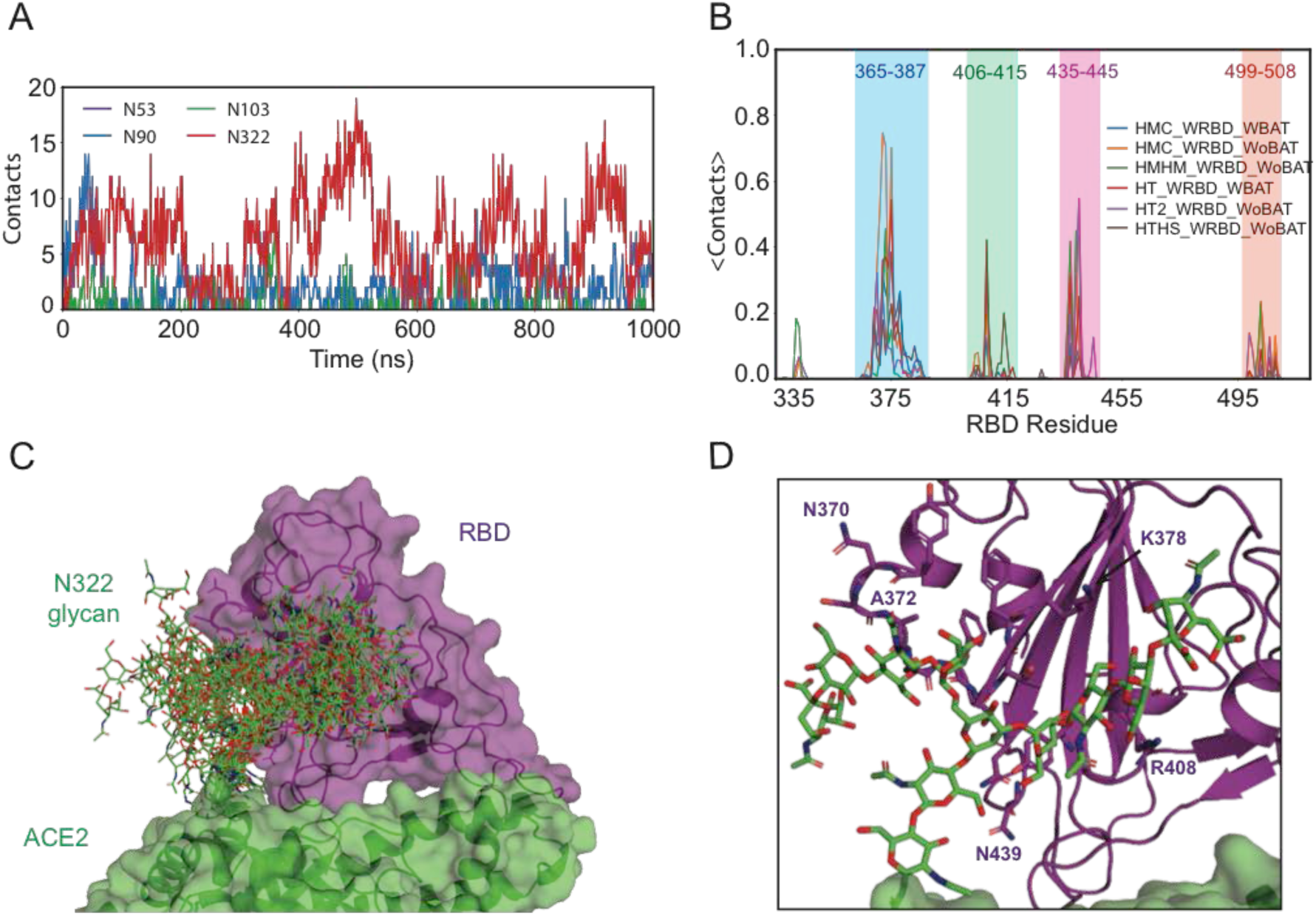
Interaction of ACE2 glycans with SARS-CoV-2 spike RBD. (*A*) Number of residue-residue contacts between the ACE2 glycans and the RBD in the HMC_WRBD_WBAT setup (see SI Appendix, Fig. S4 for all setups). (*B*) Average number of residue-residue contacts between the N322 glycans and the RBD residues. Color shading highlights the four main interaction regions. (*C*) Simulation ensemble of the N322 glycan interacting with the RBD (from the HMC_WRBD_WBAT simulation). (*D*) Close-up of the interaction between the N322 glycan and the RBD in a representative snapshot of *C*.

The N322 glycan is also a major interaction partner of the N343^RBD^ glycan on the RBD of the virus (SI Appendix, Fig. S6). The N322 glycan on ACE2 and the N343^RBD^ glycan on S have on average 1-3 interactions during the simulations.

### The N322 glycan has a specific binding site on the RBD

The finding that the N322 glycan has the highest number of interactions with the RBD prompted us to investigate whether this glycan is binding to a particular region of the RBD. To identify the structural region of the RBD interacting with the N322 glycan, we calculated the number of residue-residue interactions between glycan monosaccharides and each residue of the RBD. The resulting interaction map pinpointed regions in the RBD involving residues 365-387 and three smaller patches near residues 406-415, 435-445, and 499-508 (Fig. 2*B,C*). In these regions, the N322 glycan is interacting mainly with Y369-K378, R408, N437, N439, and V503 (Fig. 2*D*). Among these residues, the N322 glycan competes with the N90 glycan for the interaction with R408 (SI Appendix, Fig. S7). In one of the experimental ACE2-RBD structures, this residue is in direct interaction with the N90 glycan (8). Therefore, the presence of the N90 glycan interferes with N322 glycan binding to the RBD.

The ACE2-RBD binding site has a non-polar core that is surrounded by several polar and charged residues (SI Appendix, Fig. S8). The interaction of the N322 glycan with the binding site is mainly governed by weak hydrogen bonds (C-H…N/O) with 6-8 interactions per complex. Polar hydrogen bonds (3-5 interactions per complex) and hydrophobic interactions (3-5 interactions per complex) also contributed significantly to the binding affinity (SI Appendix, Table S2).

### The N90 glycan interacts weakly with the bound RBD

In 8 out of the 12 independent ACE2-RBD complexes in six different simulation setups, the N90 glycan had on average 2-5 interactions with the RBD, and at least some interactions in the remaining four simulations (SI Appendix, Fig. S4). For comparison, the two glycans at sites N53 and N103 have on average less than one interaction in all complexes. However, the N53 glycan also has occasional, transient interactions with the N343 glycan.

### The N90 glycan shields ACE2 from RBD binding

The simulations of ACE2 without the RBD allowed us to explore the possibility that the glycans near the RBD binding site might block ACE2-RBD binding. First, we calculated the number of interactions between the glycan at each glycosylation site and the residues involved in RBD binding, as identified in the five simulations of ACE2 in the absence of the RBD. Note that in each simulation we had two copies of ACE2, giving us altogether 5×2=10 copies to analyze. Of all glycans on ACE2, the one at the N90 position interacts most strongly with the uncomplexed RBD binding site (Fig. 3*A* and S9). The N90 glycan has on average 1-3 interactions with the RBD in most of the simulations (in 7 out of the 10 independent RBD-ACE2 complexes). Three complexes have less than one interaction on average. In the two complexes with the lowest number of interactions, inter-glycan interactions with the N322 glycan interfere with the interactions between the N90 glycan and the RBD binding site (SI Appendix, Fig. S10).

**Figure 3.**
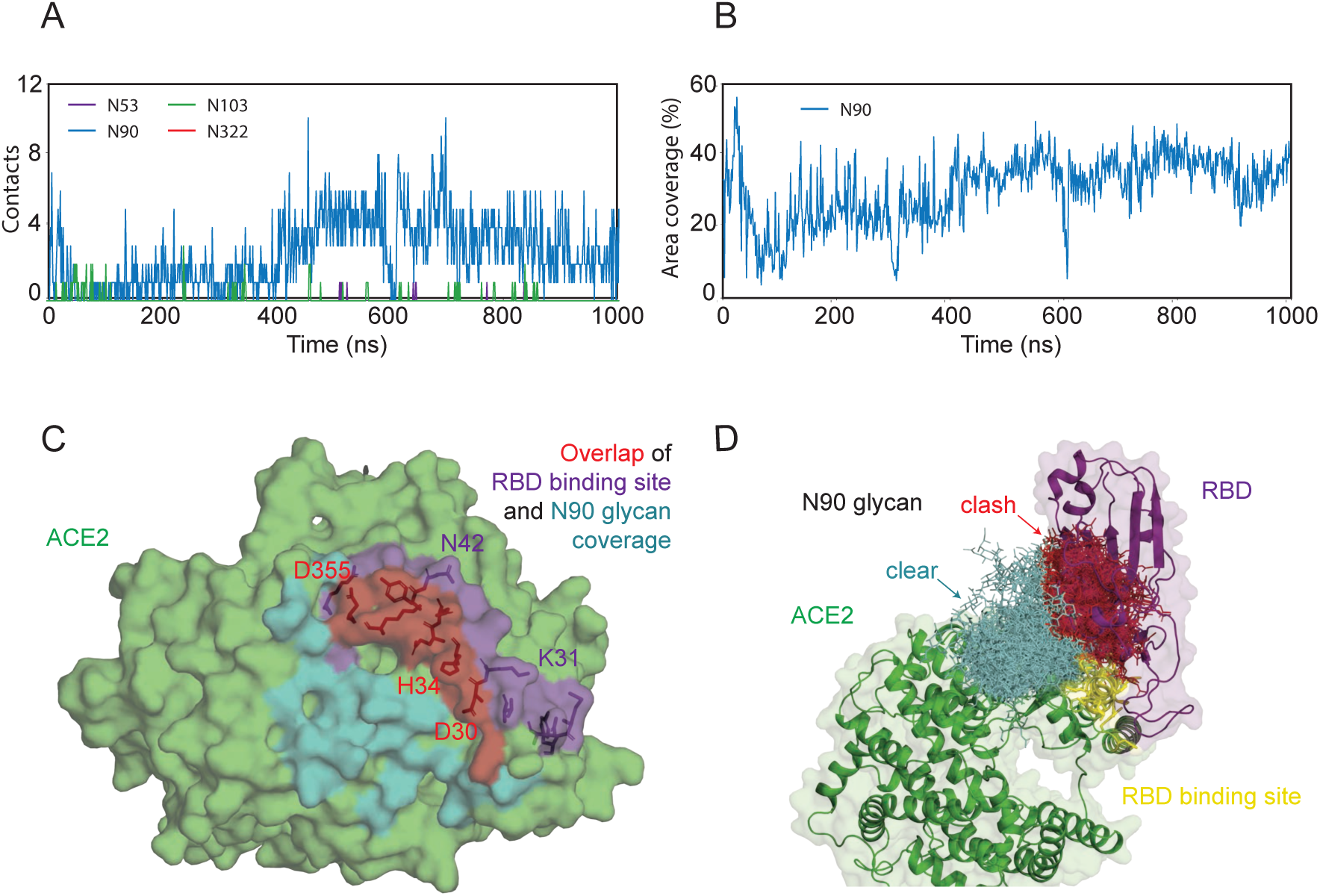
Interaction of ACE2 glycans with RBD binding site. (*A*) Number of residue-residue contacts between the ACE2 glycans and the RBD binding site in the absence of the RBD in the HMC_WoRBD_WBAT setup (see SI Appendix, Fig. S9 for all setups). (*B*) Fraction of RBD binding-site area covered by the N90 glycan from SASA calculations using a 5-Å probe for the HMC_WoRBD_WBAT setup. Results for all setups using different probe sizes are shown in SI Appendix, Figs. S12-S14. (*C*) RBD binding site shielded by the N90 glycan. The RBD binding site, the area shielded by the N90 glycan, and their overlap are colored purple, cyan, and red, respectively. (*D*) Ensemble of the N90 glycan during the simulation interacting with the RBD binding site in the HMC_WoRBD_WBAT setup. Steric clashes between glycan and RBD are illustrated by superimposing the RBD according to the HMC_WRBD_WBAT simulation. The RBD binding site is colored yellow. The glycans are shown in sticks. Glycans clashing with the RBD are colored red and those without clashes are colored cyan.

Second, we compared the solvent-accessible-surface area (SASA) of the binding region in ACE2 in the presence and absence of each glycan. Using different probe sizes allowed us to investigate distinct types of blocking. Smaller probes (1.4 Å) detect the regions in the binding site that are in a direct contact with the glycan, whereas larger probes identify regions shielded by the glycan without direct interactions and they are a better measure to check the accessibility of large molecules such as antibodies to the region. The glycans are covering on average 6.1% (3.8-12.2%), 25.6% (16.3-32.0%), and 41.4% (32.1-54.6%) of the binding site based on SASA calculations using probe sizes of 1.4, 5, and 10 Å, respectively (SI Appendix, Fig. S11). The N90 glycan is the main glycan blocking the binding site, covering 4.6% (2-8.5%), 19.1 (5.5-31.9%), and 27.4 (13.6-45.0%), respectively, for the three probe sizes (Fig. 3*B,C* and S12-14).

The N322 glycan has a lower number of interactions with the binding site and does not cover the binding site according to the SASA calculations. The only exception is the simulation with heterogonous glycan 1 (HT_WoRBD_WoBAT system), in which the N322 glycan covers a patch of the binding site comprised of residues E37, Y41, K353, and D355.

Considering the potential overlap between the glycans and the bound RBD, we noticed that in most of the 10 uncomplexed ACE2 molecules, the N90 glycan has a considerable overlap with a superimposed RBD, especially near the binding interface (Fig. 3*D* and S15). We conclude from this overlap that the N90 glycan interferes with RBD binding by blocking the interface.

## Discussion

Viral and human proteins exposed at the outer surface of virions and cells, such as SARS-CoV-2 spike and human ACE2, are heavily glycosylated. However, on reconstituted proteins used for in-vitro experiments, glycosylation is typically lacking entirely or not matching the in-situ pattern. Therefore, without detailed chemical and structural knowledge of the in-situ glycan coat, the effect of glycosylation on complex formation is often ignored in modeling. MD simulations allow us to address this challenge and to explore the effects of a variable glycan coat on protein-protein interactions.

Here we performed MD simulations of the fully glycosylated ACE2 receptor bound to the RBD of SARS-CoV-2 spike as well as unbound. These simulations gave us a detailed picture of the role of ACE2 glycans in the binding of the SARS-CoV-2 spike protein. The simulations showed contrasting effects of ACE2 glycosylation, weakening the binding of SARS-CoV-2 spike in case of the N90 glycan, strengthening the binding in case of the N322 glycan, and neutral in case of other sites.

### N90 glycosylation protects against infection

The protective effect of the N90 glycan seen in our simulations is consistent with reports in the literature on infectivity dependence on human genetic variants. Studies after the SARS outbreak in 2003 showed that the N90 glycosylation might reduce infectivity (17). A recent deep mutational analysis demonstrated that any mutation in N90 and T92 is increasing the binding affinity for the SARS-CoV-2 spike protein (18). Based on an analysis of human ACE2 polymorphism, T92I is among the human ACE2 variants that were predicted to increase susceptibility (19). ACE2 sequence comparison among species shows that several species including ferret, civet, and pig without an N90 glycosylation site can bind efficiently to the spike protein (SI Appendix, Fig. S1). Interestingly, if four residues at the N90 glycosylation site (residue 90-93) from the civet sequence are introduced into human ACE2, the SARS-CoV spike protein binds substantially more efficiently to the human receptor (17).

### N322 glycosylation likely aids infection

By contrast, the N322 glycan is crucial for RBD binding. In fact, almost all mutations removing this glycosylation site are detrimental to the binding of the RBD (18). Interestingly, mouse and rat are among the species where SARS-CoV-2 and SARS-CoV do not bind via the ACE2 receptor (6, 17, 20). Mutations at the murine and rat 322 position (H and Q in mouse and rat respectively) do not permit glycosylation at this site (SI Appendix, Fig. S1). The only exception is pangolin, where the ACE2 receptor can efficiently bind to the spike protein (20) whereas the N322 position is mutated and cannot be glycosylated.

Our MD simulations show that the N322 glycan is binding into a well-defined region in the RBD with a core of the residues 369-378. Comparison of the RBD sequences among different CoV viruses shows that the identified binding site is mostly conserved (Fig. 4*A*).

**Figure 4.**
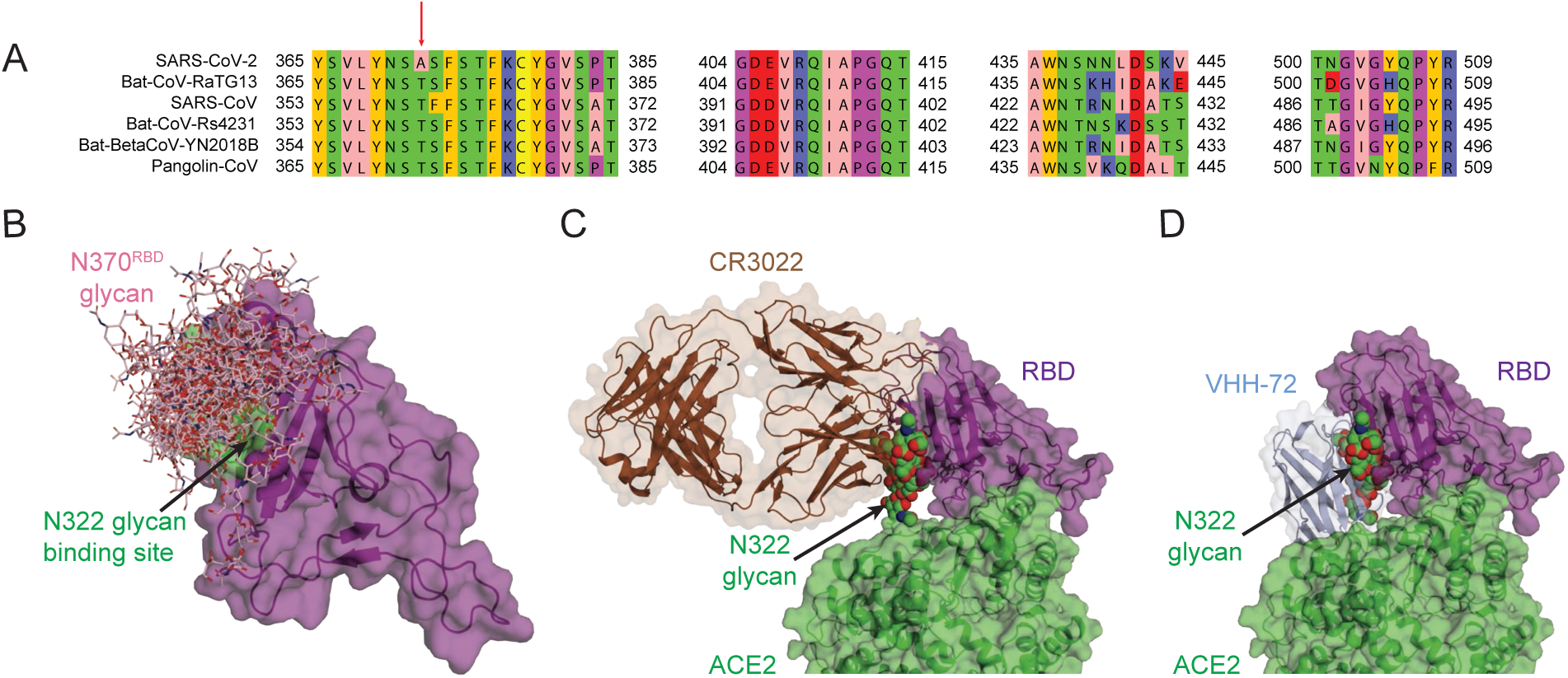
N322 binding site in RBD. (*A*) Sequence alignment of the N322 binding site in different coronaviruses. The red arrow indicates the difference between SARS-CoV and SARS-CoV-2 in the glycosylation motif of the N370^RBD^ site. (*B*) Ensemble of N370^RBD^ interacting with the N322 binding site (green) in the simulation of the RBD alone. (*C*) Superimposition of antibody CR3022 (brown) targeting the cryptic epitope with a representative snapshot of the N322 glycan (space filling, green). (*D*) Superimposition of antibody VHH-72 (light blue) targeting the region near the cryptic epitope with a representative snapshot of the N322 glycan (space filling, green).

### Lack of N370 glycosylation opens a window onto spike

We also explored the effect of spike glycosylation on ACE2 binding. Indeed, antigen glycosylation governs host immune responses. In the case of SARS-CoV-2, the glycans are covering almost the entire surface of the spike protein, suggesting that the virus can avoid the host immune system in a stealth fashion. In this sense, the absence of the N370 glycosylation site in the RBD of SARS-CoV-2 compared to SARS seems perplexing, as this mutation leaves a patch of the RBD exposed as a target for host immunological responses (Fig. 4*A*). Simulating the RBD^CoV2^ with a glycan at N370 shows that the N370^RBD^ glycan interacts with the same core of residues (Y369-K378) as the N322 glycan (Fig. 4*B*).

### N322 glycan locks into cryptic binding site targeted by neutralizing antibodies

The site on the RBD binding the N322 glycan, as seen in our simulations, is a prime target for neutralizing antibodies. First, this region contributes positively to the binding of the spike protein and, second, in contrast to the SARS-CoV, this region is not covered by a glycan and presumably exposed to the solvent (at least in some conformations of the spike protein). The recent hunt for neutralizing antibodies against the SARS-CoV-2 spike protein revealed a presumably cryptic epitope in the RBD targeted by a class of antibodies. Remarkably, this cryptic epitope is the same region that our MD simulations revealed as the binding site for the N322 glycan. An antibody (CR3022), which has been obtained from a convalescent SARS-CoV infected patient, has a binding site that overlaps significantly with that of the N322 glycan in our simulations (Fig. 4*C*) (21). Intriguingly, the lower affinity of RBD^CoV2^ for CR3022 is associated with the absence of the N370^RBD^ glycan. The VHH-72 antibody developed for RBD^CoV^ has a similar binding site as CR3022 (Fig. 4*D*) (22). This antibody is mainly interacting with Y356^CoV^ (Y369^CoV2^), S358^CoV^ (S371^CoV2^), K365^CoV^ (K378^CoV2^), C366^CoV^ (C379^CoV2^), R426^CoV^ (N439^CoV2^), and Y494^CoV^ (Y508^CoV2^), most of which interact with the N322 glycan in the RBD^CoV2^. Both antibodies have a suboptimal affinity for RBD^CoV2^, as they were raised against RBD^CoV^. This shows the need for developing neutralizing antibodies targeting specifically RBD^CoV2^. Therefore, details of the interactions between the N322 glycan and this epitope are central for a rational design of antibodies.

In conclusion, we established a molecular picture for the role of ACE2 glycosylation in the binding of the SARS-CoV-2 that provides mechanistic insights. We hope that our findings can serve as a basis for the rational development of neutralizing antibodies and small molecules that target the N322-glycan binding site in the RBD of the SARS-CoV-2 spike protein, and possibly of small molecules that mimic the protective effect of the N90-glycosylation variant on human ACE2.

## Materials and Methods

### The ACE2 complex

The coordinates of the ACE2 complex were taken from PDB ID: 6M17(9) re-refined by Tristan Croll. The complex contains a dimer of the ACE2 receptor in complex with the RBD and also the B^0^AT1 transporter. The ACE2 receptor was simulated in the dimeric form and in the absence or presence of the RBD and also the B^0^AT1 transporter in different simulation setups (See SI Appendix, Table S1).

### Glycosylation

Based on the cryoEM structure of the complex, the ACE2 receptor, the RBD, and the B^0^AT1 transporter contain seven (N53, N90, N103, N322, N432, N546, N690), one (N343), and five (N158, N182, N258, N354, N358) N-glycosylation sites, respectively. We considered five glycosylation patterns in the simulations (See Fig. 1 and SI Appendix, Table S1). Two zinc ions in the peptidase binding sites of ACE2 were retained during the simulations. In two sets of systems, the glycosylation sites in the ACE2 receptor and the B^0^AT1 transporter were uniformly glycosylated with either a complex N-glycan or a high mannose N-glycan (the homogenous glycosylation patterns in Fig. 1). In three sets of systems, each site in the ACE2 receptor was glycosylated with the most frequent N-glycan based on the MS data (13) (the heterogeneous glycosylation patterns in Fig. 1). In the heterogeneous glycosylated systems, an O-glycosylated site (T730) was also glycosylated based on the MS data (13) (the heterogeneous glycosylation patterns in Fig. 1). The glycan at the glycosylation site in the RBD (N343) was added based on the MS analysis of the SARS-CoV-2 spike protein (14).

### System setup

The interaction of the ACE2 receptor with the RBD of the spike protein was studied with all-atom explicit solvent MD simulation using GROMACS v2019.6 (23). The lipid bilayers of palmitoyl oleoyl phosphatidyl-choline (POPC), palmitoyl oleoyl phosphatidyl-ethanolamine (POPE), palmitoyl oleoyl phosphatidyl-serin (POPS), palmitoylsphingomyelin (PSM), and cholesterol lipids were created using the CHARMM-GUI webserver (24). Lipid ratios are listed in SI Appendix, Table S3. These systems were solvated with water and 150 mM NaCl, resulting in boxes of ∼21×21×26 nm^3^ (∼1,000,000 atoms).

After 2,000 steps of steepest descent energy minimization, the membrane patch was equilibrated first for 1 ns of MD simulation in an NVT ensemble with a 1-fs time step and then in an NPT ensemble (7.5 ns) with a 2-fs time step using a Berendsen thermostat and barostat (25). Production simulations of 1 µs each were run with a 2-fs time step at a temperature of 310 K and a pressure of 1 bar in an NPT ensemble using a Nosé-Hoover thermostat (26) and a semi-isotropic Parrinello-Rahman barostat (27) with a characteristic time of 5 ps and a compressibility of 4.5×10^−5^ bar^-1^.

The all-atom CHARMM36m force field was used for protein, lipids and ions, and TIP3P was used for water molecules (28, 29). The MD trajectories were analyzed with Visual Molecular Dynamics (VMD) (30) and MDAnalysis package (31).

### RBD glycosylation

We performed MD simulations of an isolated RBD (chain F of PDB ID 6M17) glycosylated at positions N343^RBD^ and N370^RBD^. For this system, we used the homogenous complex glycan (Fig. 1*B* left panel). The RBD was solvated with water and 150 mM NaCl, resulting in boxes of ∼12.5×12.5×12.5 nm^3^ (∼84,000 atoms). After 2,000 steps of steepest descent energy minimization, the system was equilibrated first for 0.5 ns of MD simulation in an NVT ensemble with a 1-fs time step and then in an NPT ensemble (8.5 ns) with a 2-fs time step using a Berendsen thermostat and barostat (25). Production simulations of 1 µs each were run with a 2-fs time step at a temperature of 310 K and a pressure of 1 bar in an NPT ensemble using a Nosé-Hoover thermostat (26) and an isotropic Parrinello-Rahman barostat (27) with a characteristic time of 5 ps and a compressibility of 4.5×10^−5^ bar^-1^.

### Analysis

#### Residue-residue contacts

A pair of residues (θ_*i*_ and θ_*j*_) was considered being in contact when at least one pair *i* and *j* of heavy atoms belonging to θ_*i*_ and θ_*j*_, respectively, was within 3.5 Å in distance. For a hydrogen bond, two geometrical conditions had to be fulfilled: 1) The D-H…A distance had to be lower than 3.4 Å and 2) the D-H…A angle had to be higher than 120°, with D and H the donor and acceptor atoms, respectively. For a hydrophobic interaction, two C or S atoms had to be within 4 Å of each other.

#### Fraction of native contacts

Following Best et al. (32), two heavy atoms *i* and *j* are considered to form a native contact if their distance *r*_*ij*_^0^ in the cryoEM structure is less than 4.5 Å. We then defined the fraction of native contacts *Q*(*X*) in a configuration *X* as

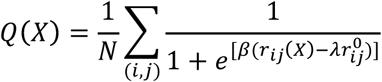

where the sum runs over the *N* pairs of native contacts (*i,j*) and *r*_*ij*_(*X*) is the distance between *i* and *j* in configuration *X*. We set the smoothing and padding parameters to β=5 Å^-1^ and λ=1.8, respectively.

#### Solvent accessible surface area

The SASA was calculated for each atom. We extended its radius by the size of the probe and determined the area of the sphere exposed to solvent. Three different probe sizes of 1.4, 5, and 10 Å were used to measure the SASA.

### Sequence alignment

The sequences of the ACE2 receptor from various species and also the different coronaviruses were aligned using T-COFFEE program (33).

## Author contributions

A.R.M. and G.H. designed research; A.R.M. performed research; A.R.M. and G.H. analyzed data; and A.R.M. and G.H. wrote the paper.

## Acknowledgments

We thank Mateusz Sikora, Florian Blanc, and Sören von Bülow for their help during system setup and analysis. This work was supported by the Max Planck Society and the German Research Foundation (SFB 807 – Membrane Transport and Communication).

---

**Figure S1.**
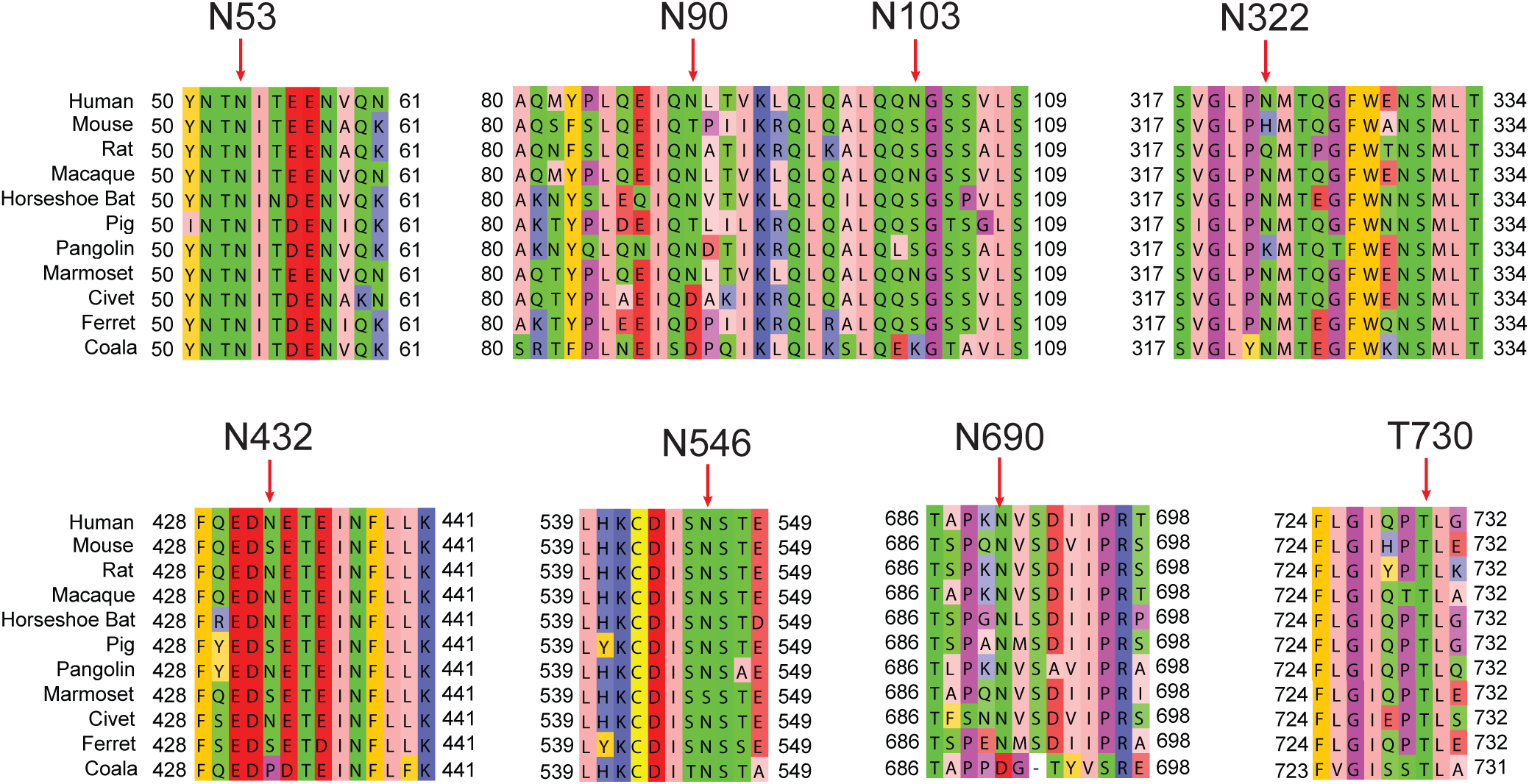
Amino acid sequence alignment of the ACE2 receptor (glycosylation sites) in human and selected species. The red arrows indicate the glycosylation sites of human ACE2 and the homologous sites in other species.

**Figure S2.**
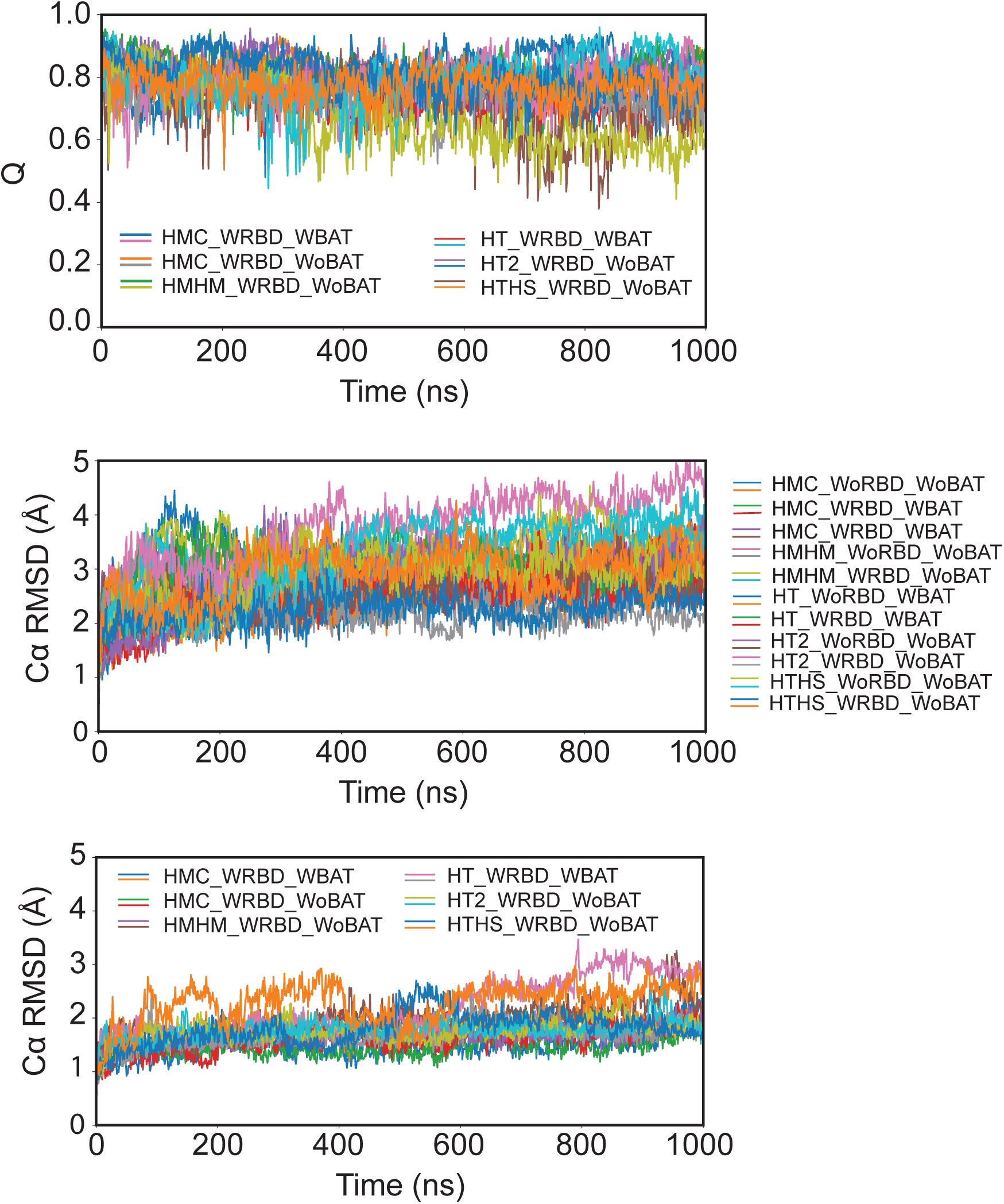
Structural and conformational stability of the ACE2 complex. Top panel shows the fraction of native contacts (Q) for the binding interface between ACE2 and the RBD during the simulation time in different simulation setups. Middle panel shows the Cα RMSD of the peptidase domains of the ACE2. The peptidase domains (residue 21-610) were first superimposed into the Cryo-EM structure using the Cα atoms and then the RMSDs were calculated. Bottom panel shows the Cα RMSD of the RBDs. The RBDs (residue 336-569) were first superimposed into the Cryo-EM structure using the Cα atoms and then the RMSDs were calculated.

**Figure S3.**
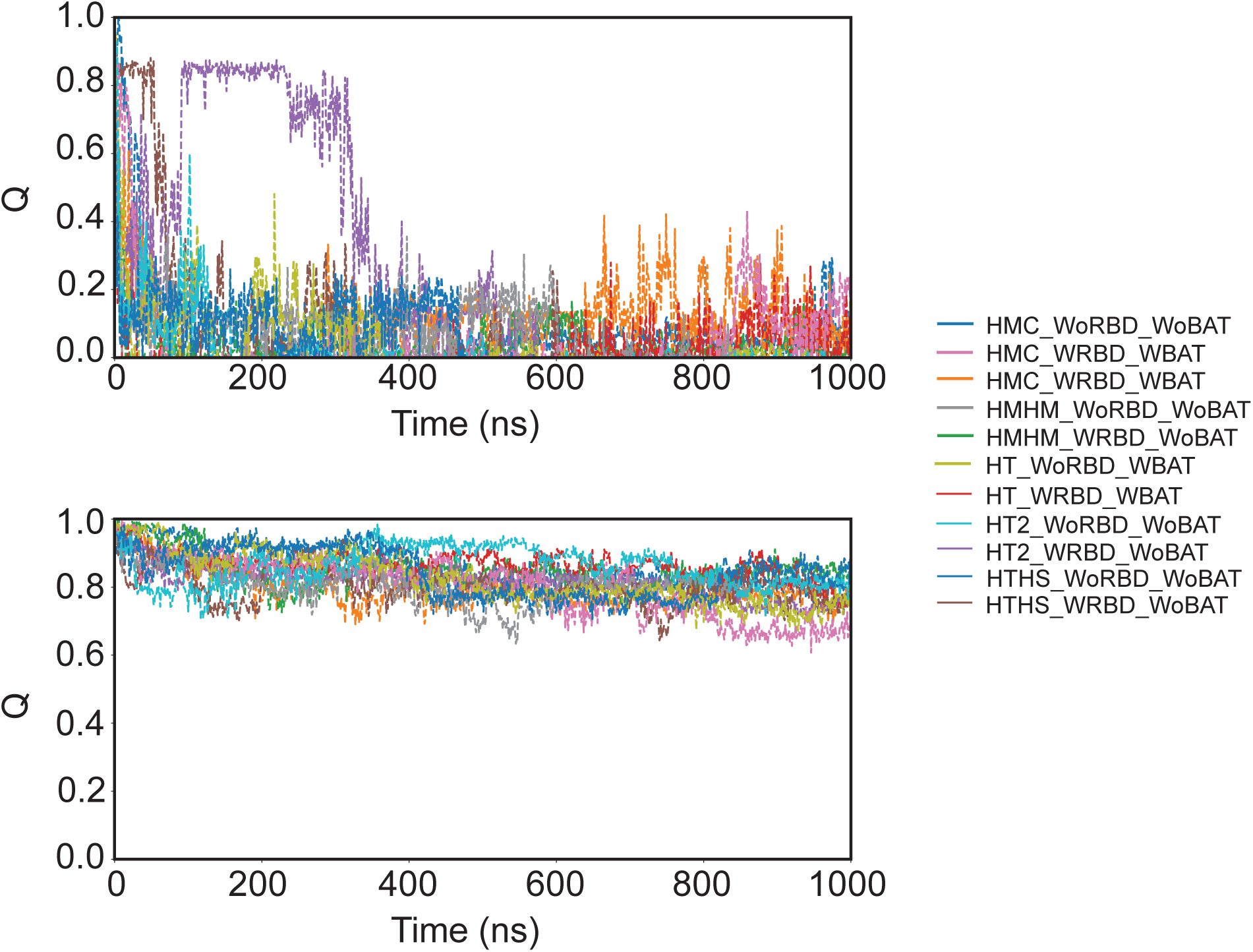
Conformational stability of the ACE2 dimerization interfaces. The fraction of native contacts (Q) is shown between two ACE2 monomers at the lower (residues 610-725) and upper interfaces (residues 110-195) during the simulation time in different simulation setups (top and bottom panels respectively).

**Figure S4.**
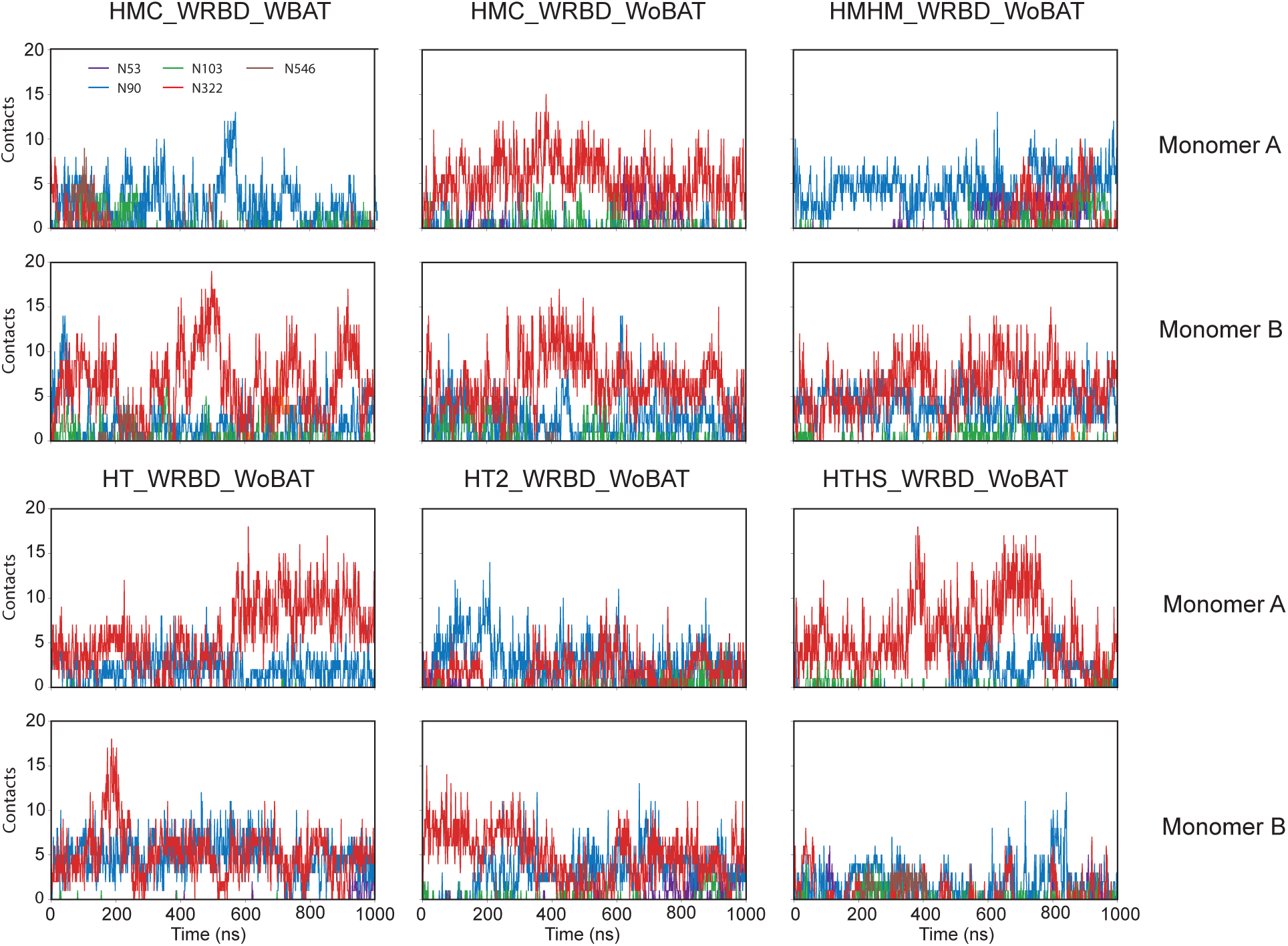
Interaction between the ACE2 glycans and the RBD. Number of residue-residue contacts is shown for all seven glycosylation sites for different glycosylation patterns (See Table S2).

**Figure S5.**
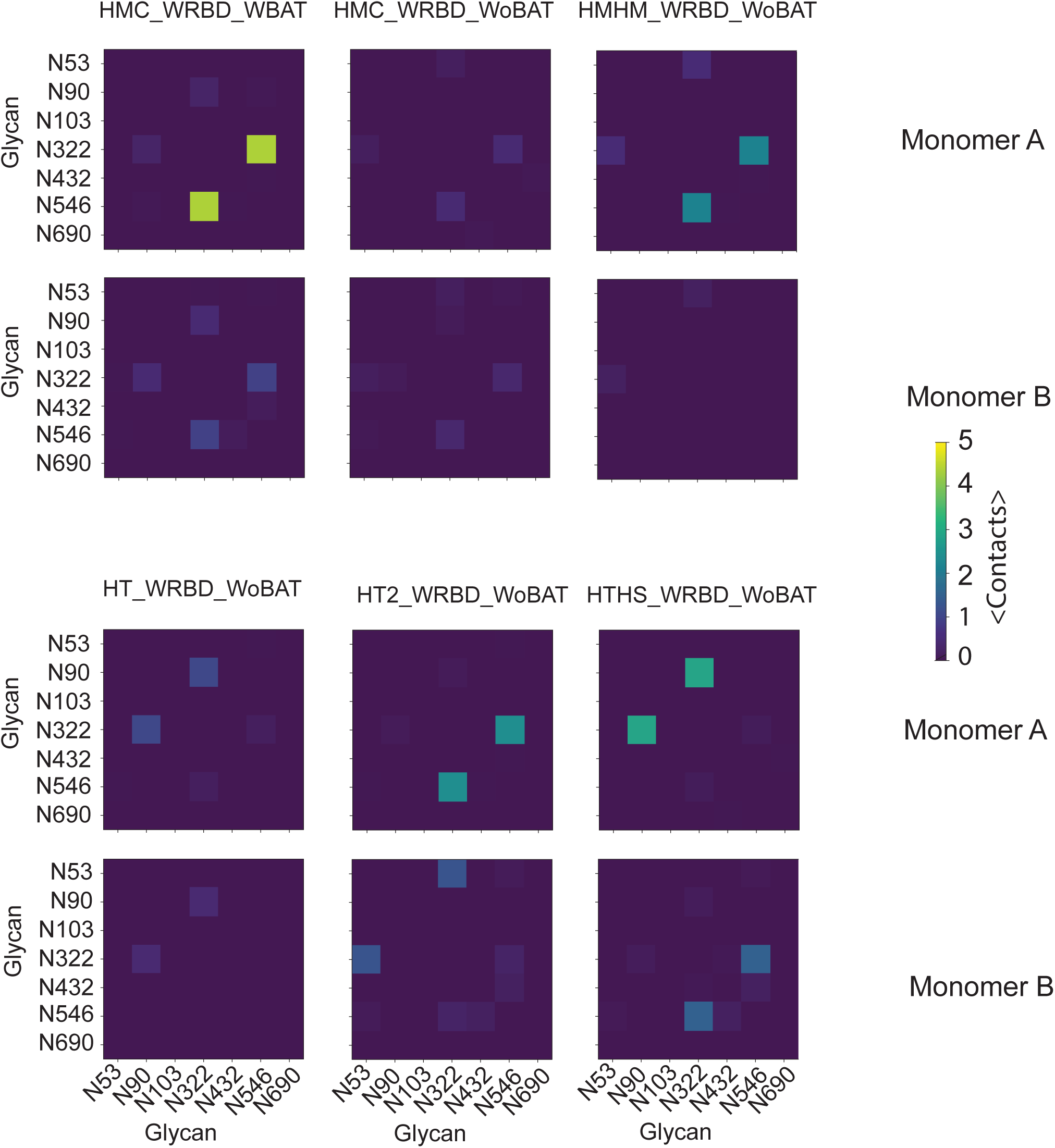
Inter-glycan interaction in ACE2-RBD complex. Average number of residue-residue contacts is shown for all glycan pairs in ACE2 for different glycosylation patterns (See Table S2) for those simulations performed in the presence of the RBD.

**Figure S6.**
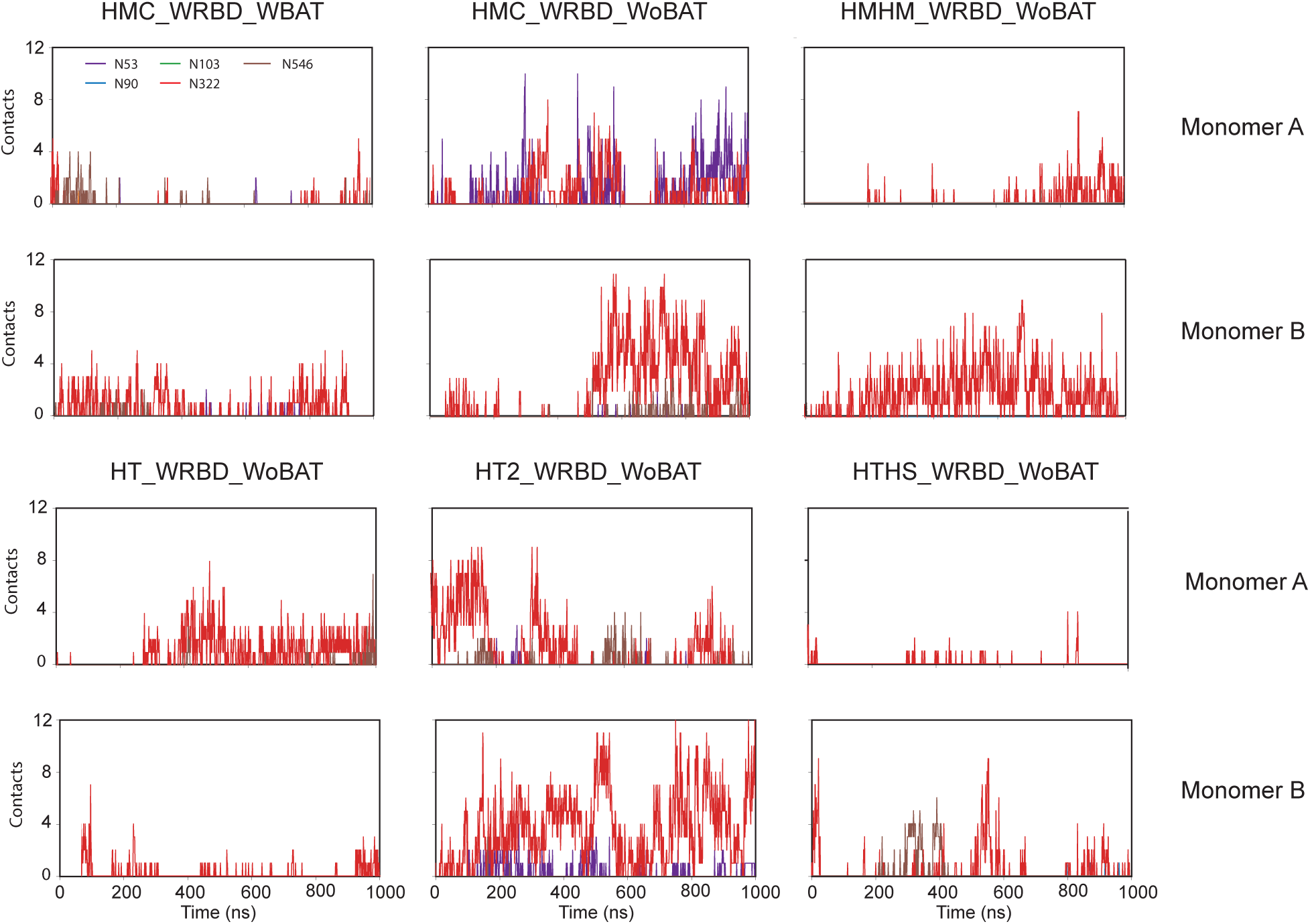
Interaction between the ACE2 glycans and the N343 glycan of the RBD. Number of residue-residue contacts is shown between 4 glycosylation sites in ACE2 near the RBD and the N343RBD glycan for different glycosylation patterns (See Table S2).

**Figure S7.**
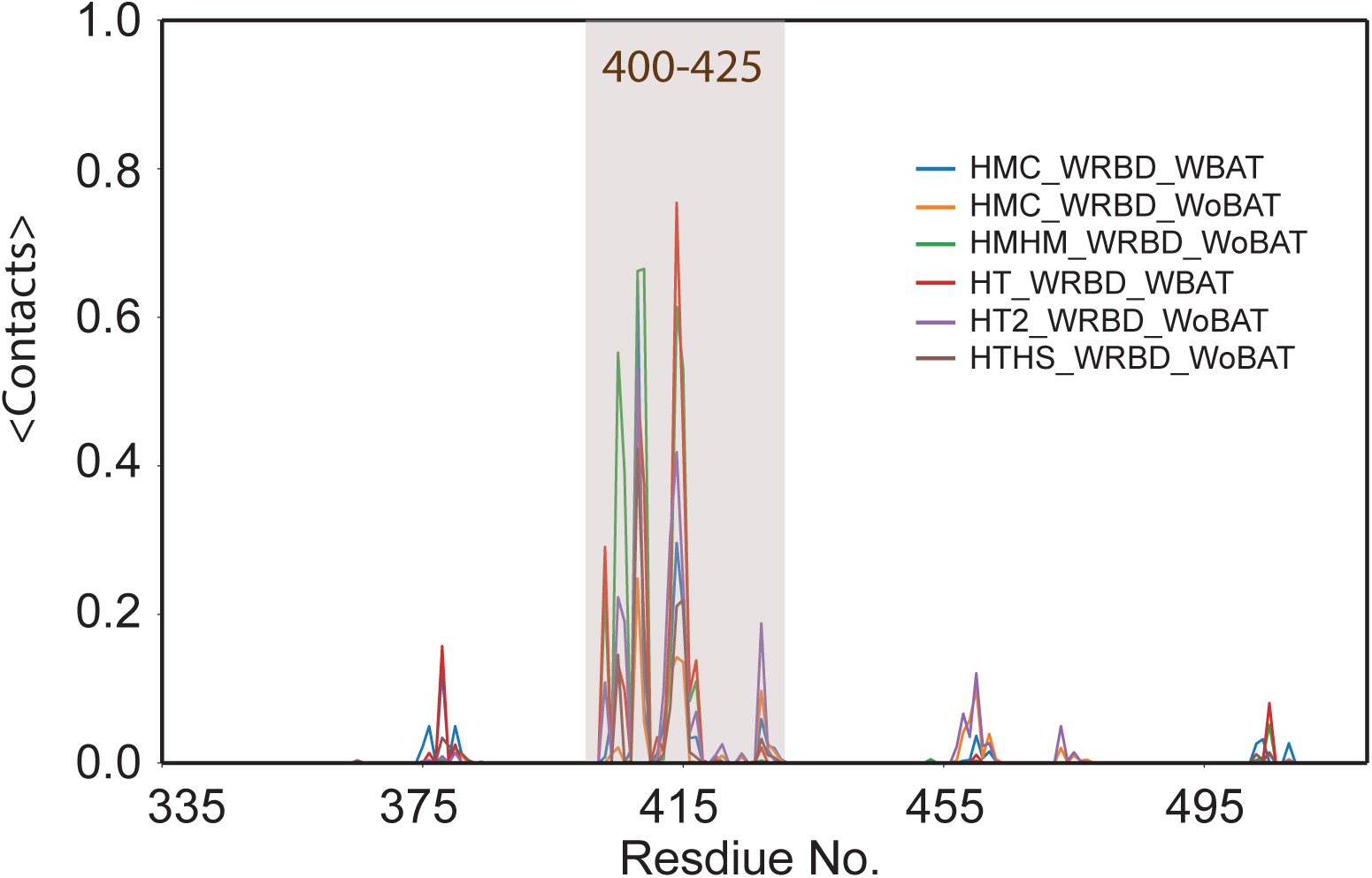
Interaction between the N90 glycan and the RBD. Average number of residue-residue contacts is shown between the N90 glycan and the RBD residues. The main interaction region is highlighted.

**Figure S8.**
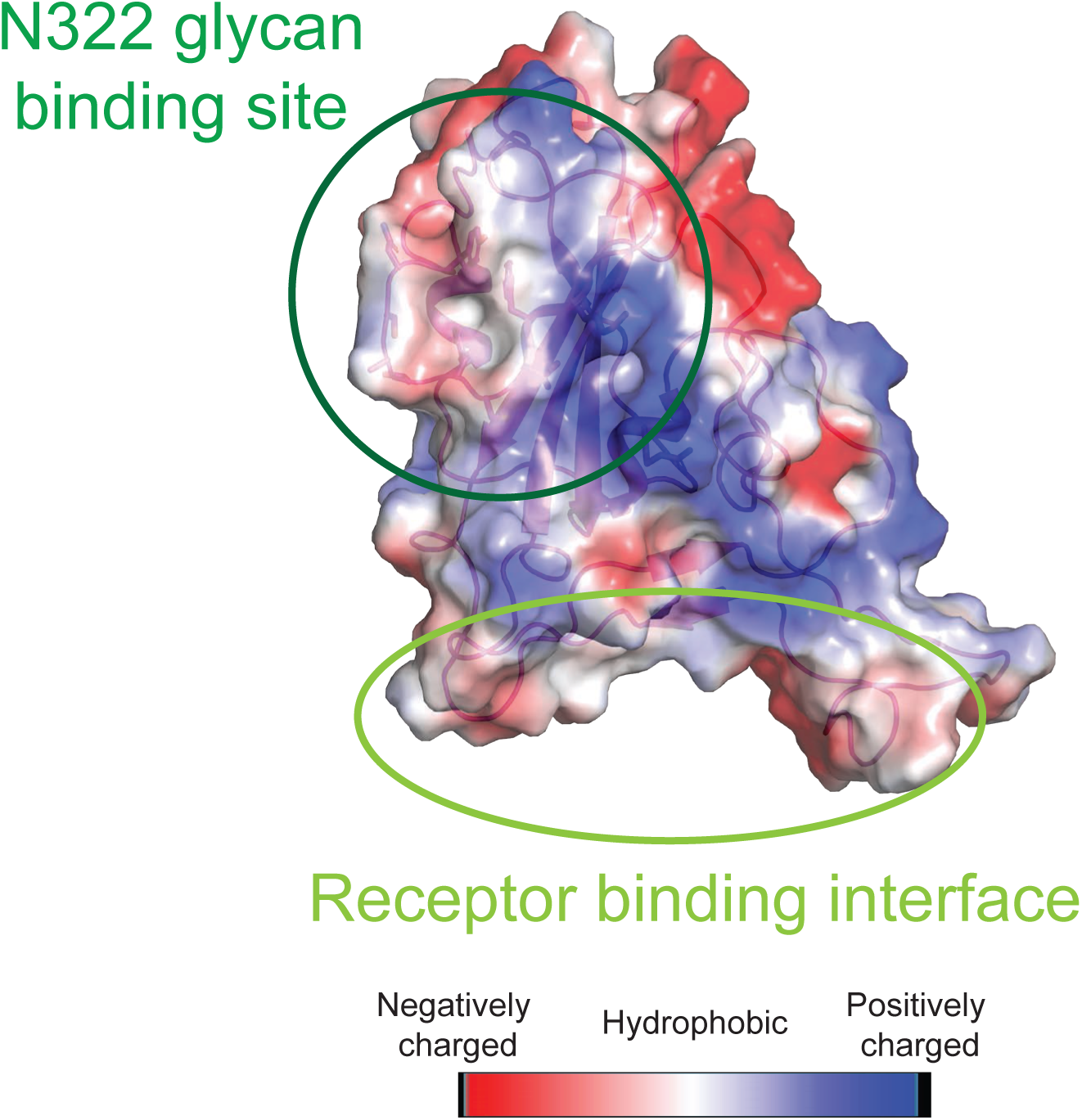
Electrostatic surface charge of the RBD. The binding site of the N322 glycan and the interaction interface of ACE2 are marked as dark green and light green ovals, respectively.

**Figure S9.**
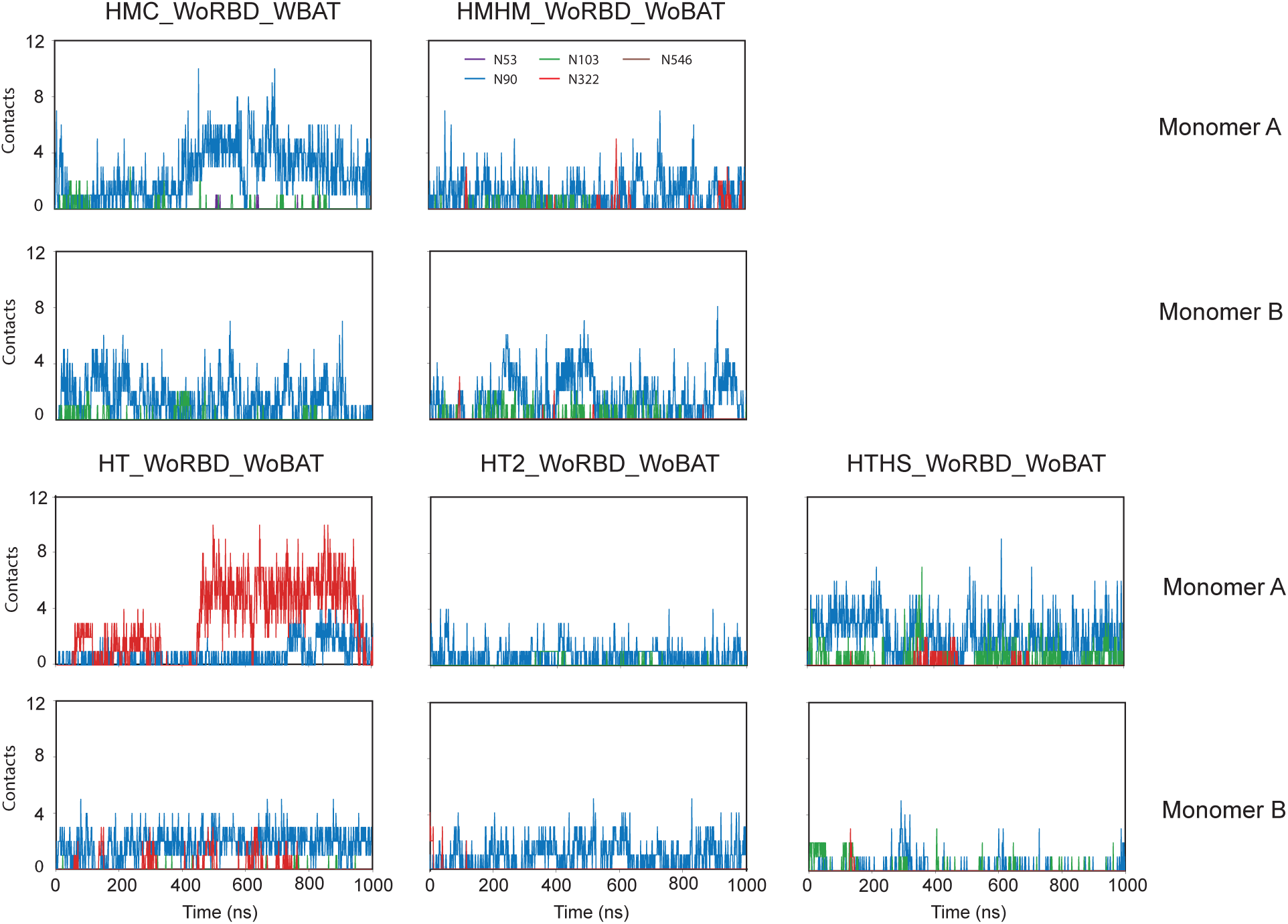
Interaction between the ACE2 glycans and the RBD binding site. Number of residue-residue contacts is shown for all seven glycosylation sites for different glycosylation patterns (See Table S2) for those simulations performed in the absence of the RBD.

**Figure S10.**
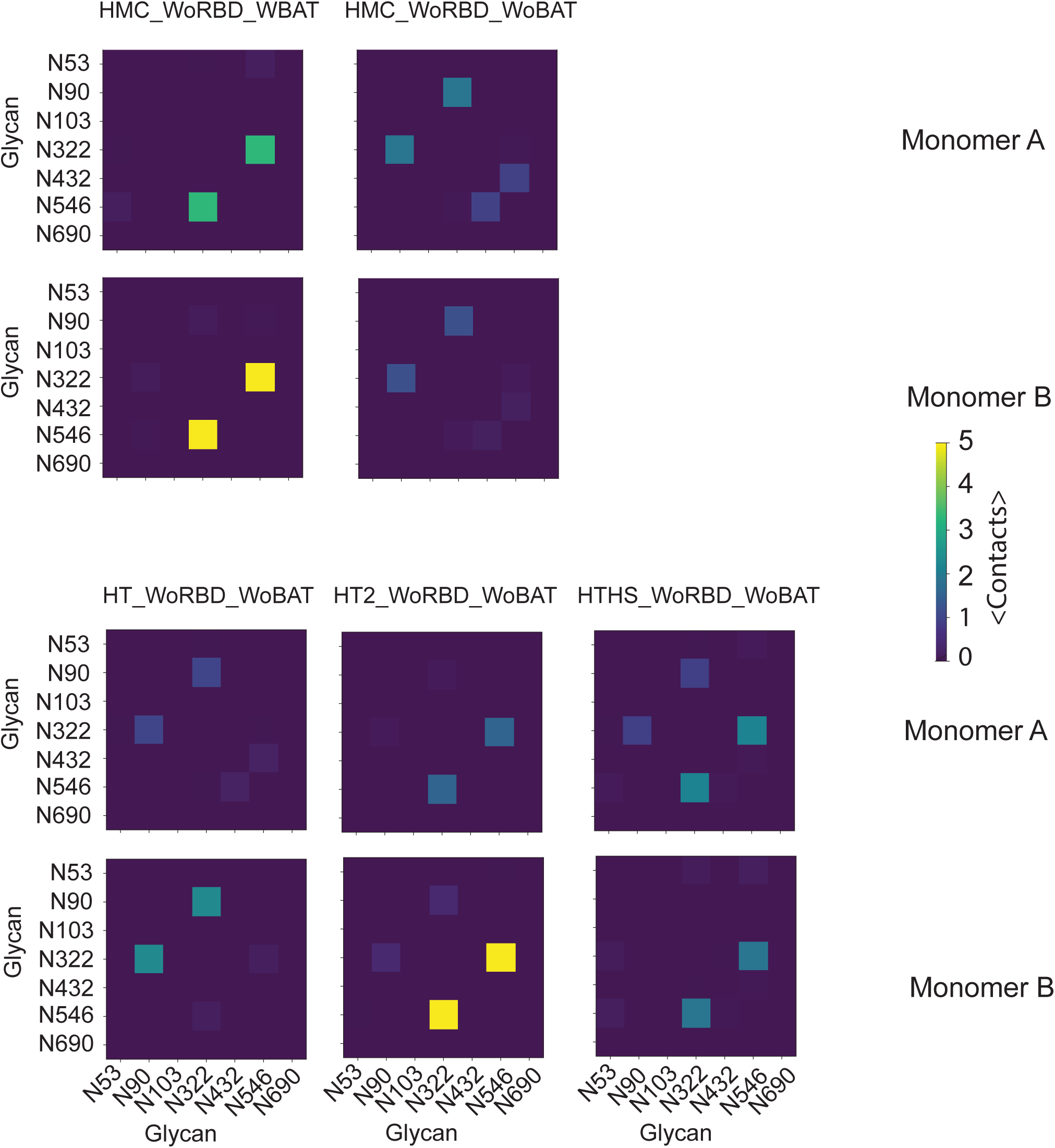
Inter-glycan interaction in the apo state of ACE2. Average number of residue-residue contacts is shown all glycan pairs in ACE2 for different glycosylation patterns (See Table S2) for those simulations performed in the absence of the RBD.

**Figure S11.**
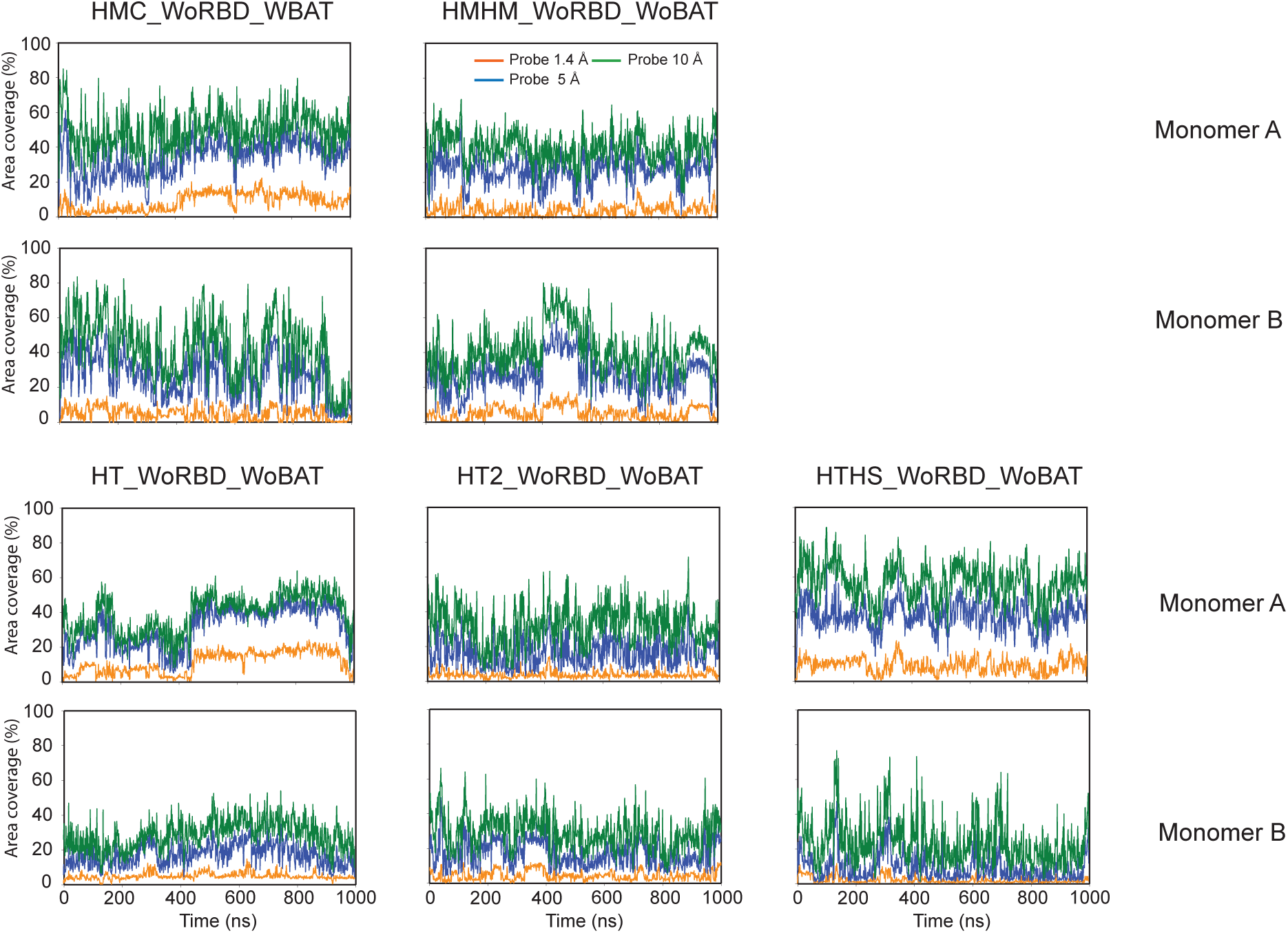
Glycan shielding of the RBD binding site. Shielding is defined as the percentage of surface covered by glycans calculated based on the difference between the SASA values of the binding site in the absence and presence of glycans. Three different probe sizes, 1.4, 5, and 10 Å, were used to calculate the SASA values.

**Figure S12.**
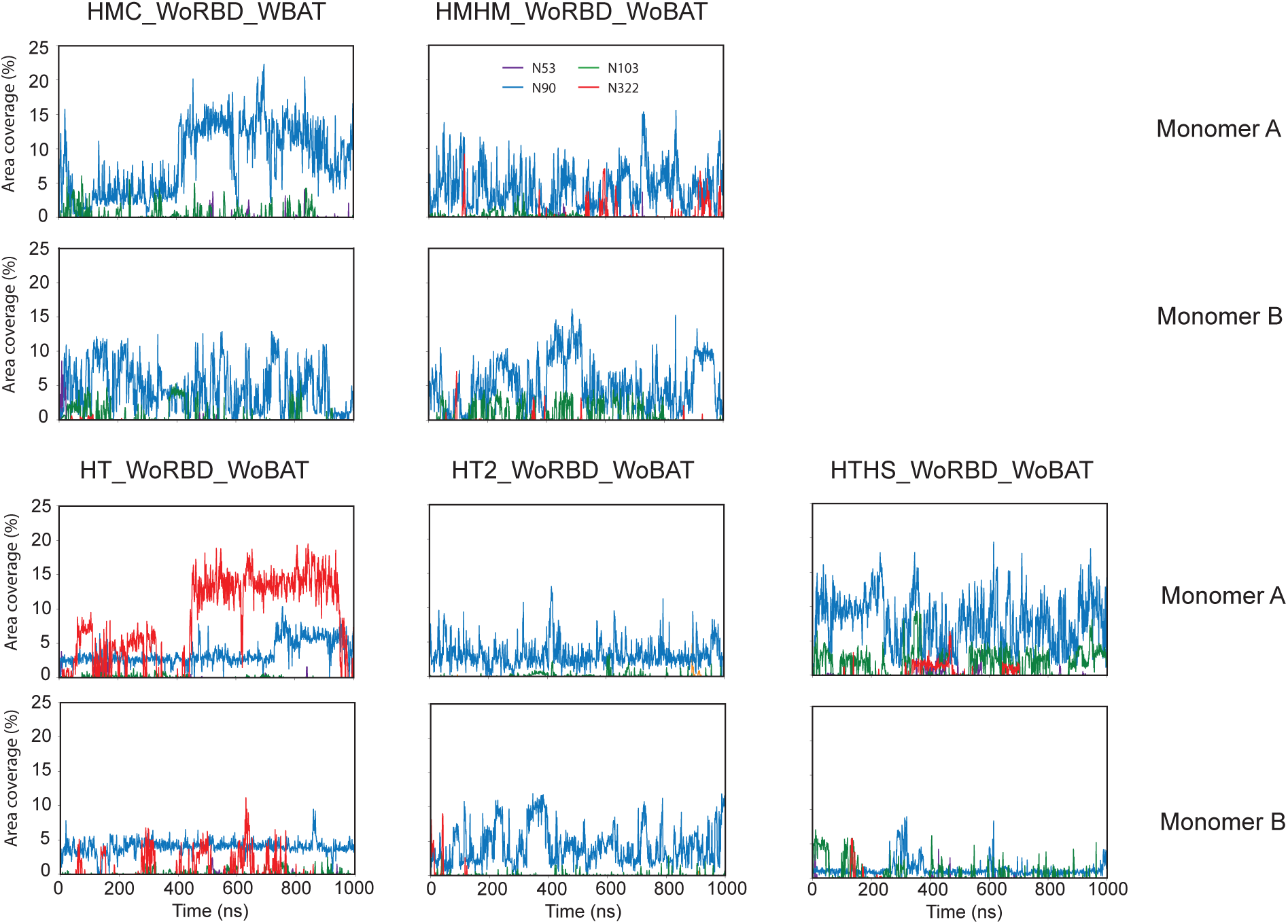
Contribution of each glycan in shielding of the RBD binding site based on a 1.4-Å probe. Shielding is defined as the percentage of surface covered calculated based on the difference between the SASA values of the binding site in the absence and presence of glycans.

**Figure S13.**
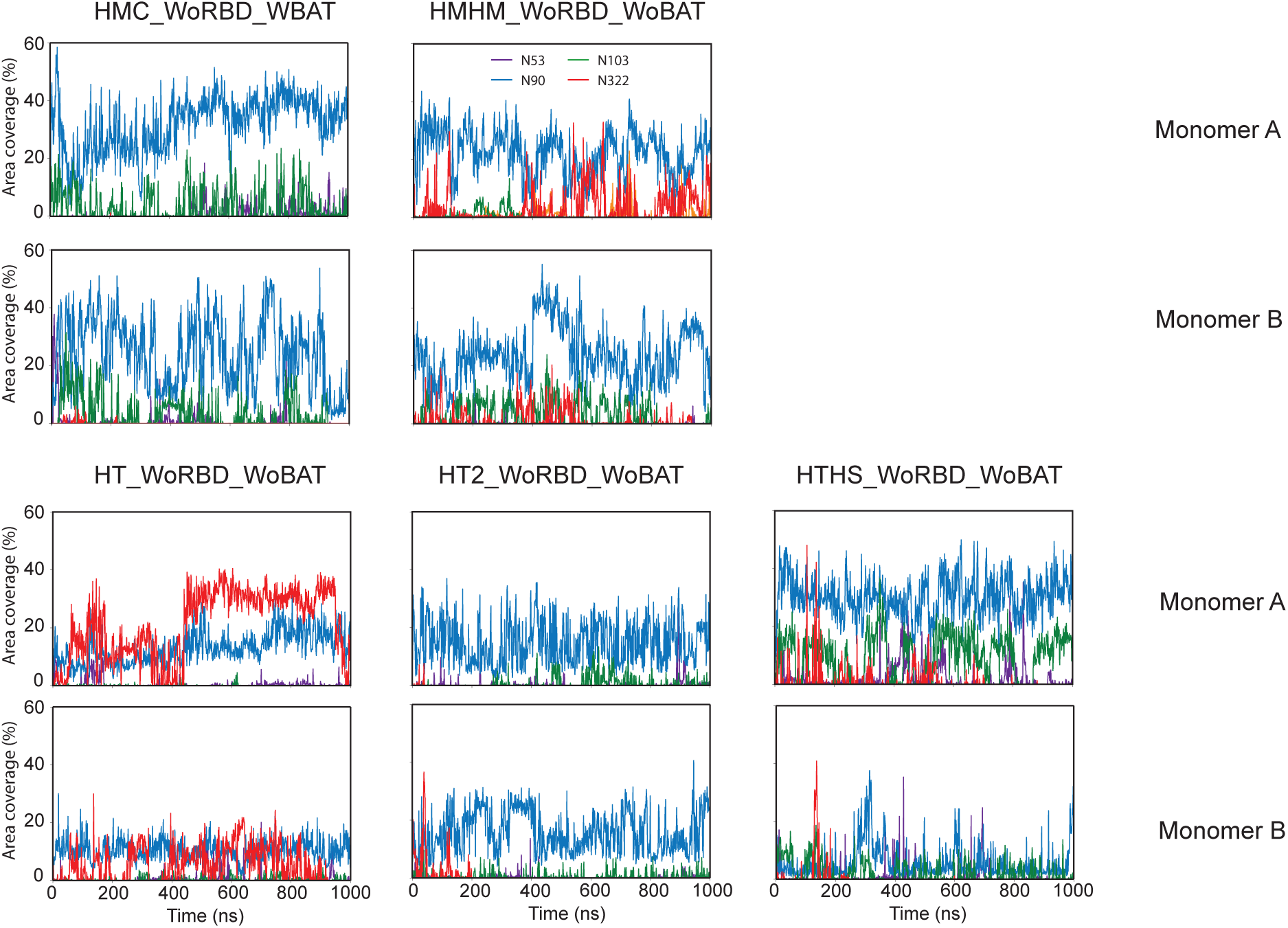
Contribution of each glycan in shielding of the RBD binding site based on a 5-Å probe.

**Figure S14.**
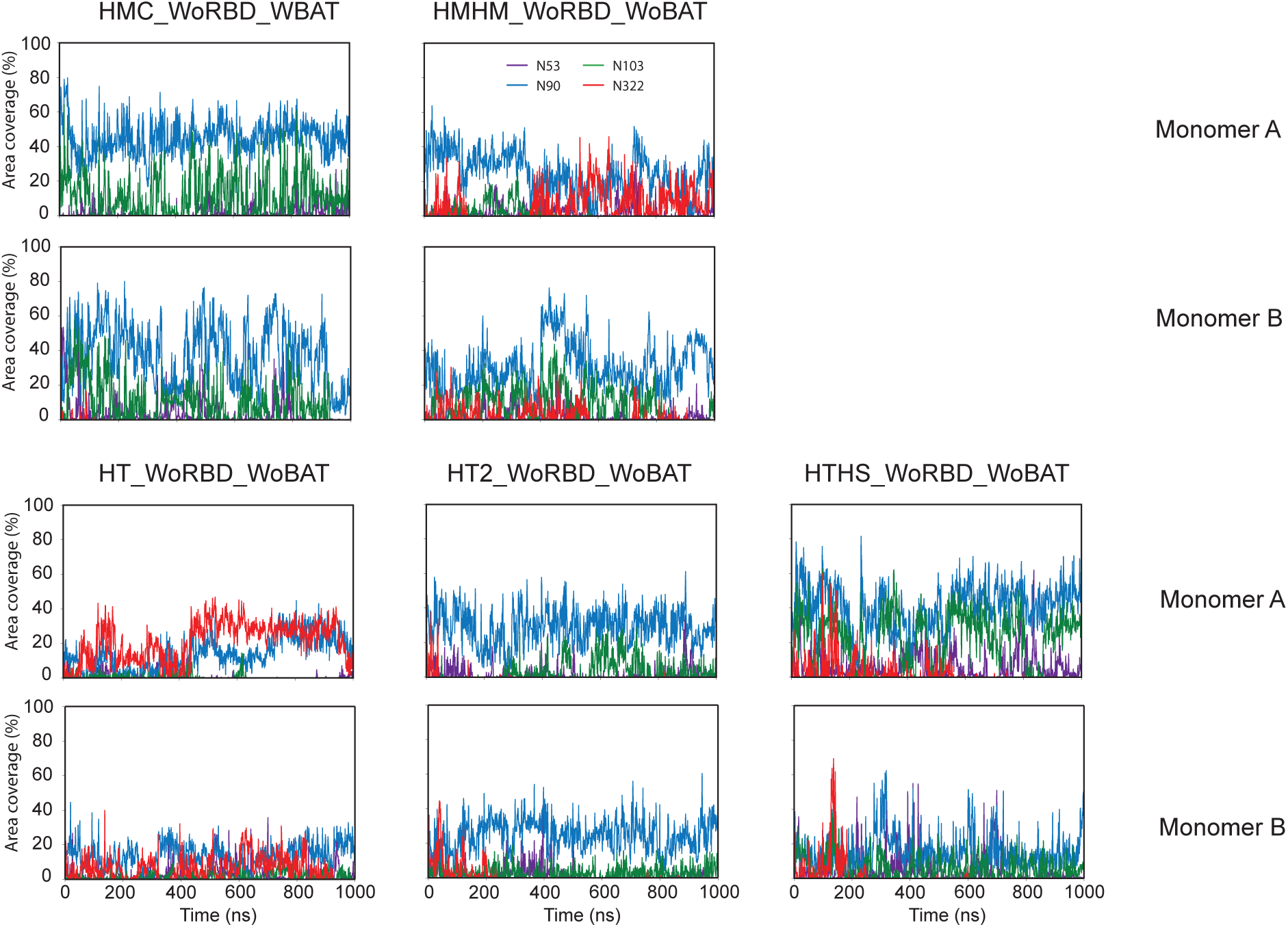
Contribution of each glycan in shielding of the RBD binding site based on a 10-Å probe.

**Figure S15.**
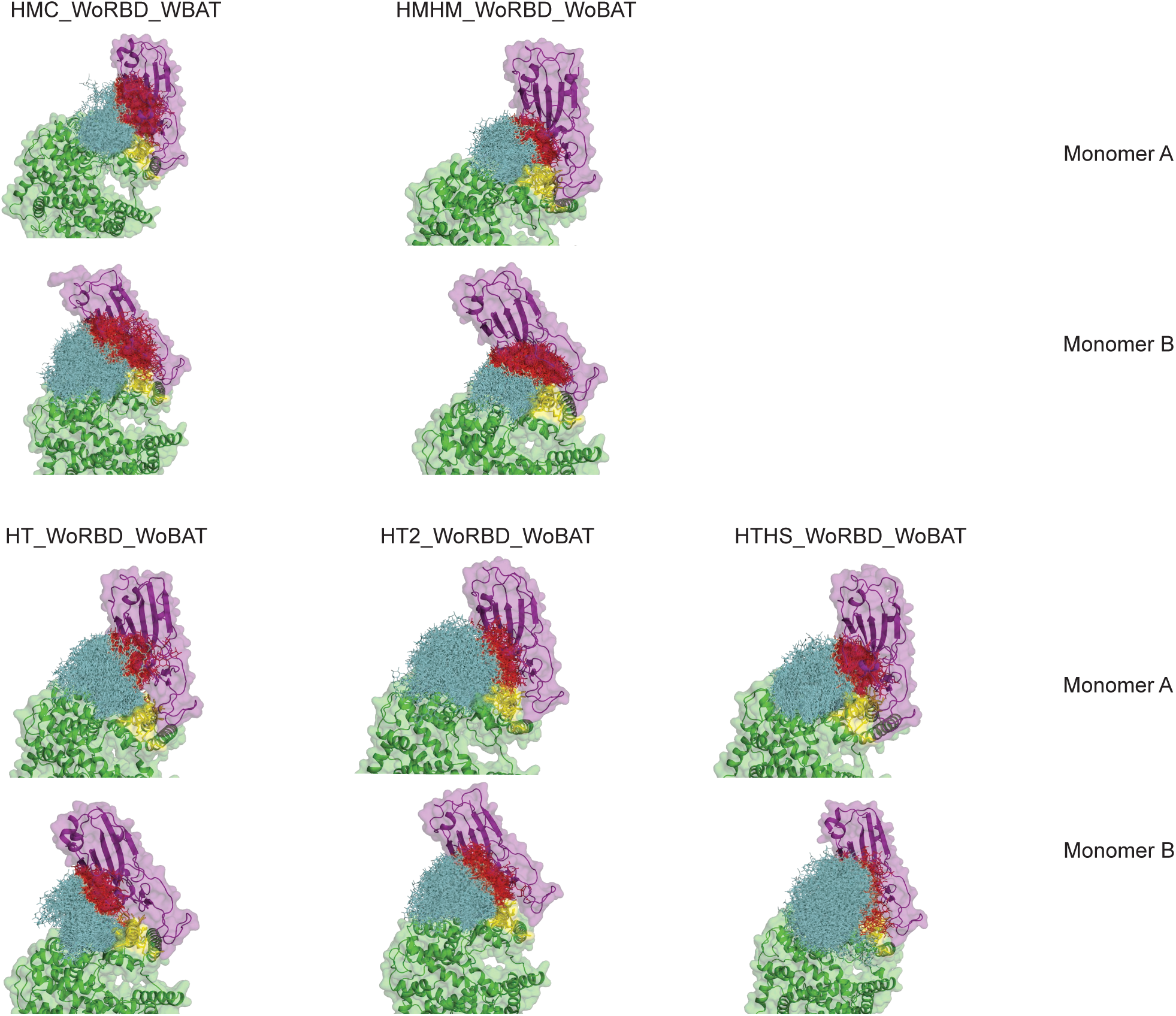
Interaction of N90 glycan with the RBD binding site during the simulations in the apo state of ACE2. An average structure of the RBD from the corresponding RBD–bound simulation is also shown in each panel. ACE2 and the RBD are colored green and purple respectively. The RBD binding site is colored yellow. The glycans are shown in sticks. Glycans that have steric clashes with the superimposed RBD are colored red and those without clashes are colored cyan.

**Table S1.** Description of the different simulations performed in this study.

| System name | Protein components |  |  | System size<br>(Å<sup>3</sup>) | Glycosylation | Length (ns) |
| --- | --- | --- | --- | --- | --- | --- |
|  | ACE | B<sup>0</sup>AT1 | RBD |  |  |  |
| HMC_WBAT_WRBD | + | + | + | 210×210×255 | Homogeneous complex N-glycan | 1000 |
| HMC_WBAT_WoRBD | + | + | - | 210×210×240 | Homogeneous complex N-glycan | 1000 |
| HMC_WoBAT_WRBD | + | - | + | 210×210×255 | Homogeneous complex N-glycan | 1000 |
| HMHM_WoBAT_WRBD | + | - | + | 210×210×255 | Homogeneous high mannose N-glycan | 1000 |
| HMHM_WoBAT_WoRBD | + | - | - | 210×210×240 | Homogeneous high mannose N-glycan | 1000 |
| HT_WoBAT_WRBD | + | - | + | 210×210×255 | Heterogeneous N-glycan 1 | 1000 |
| HT_WoBAT_WoRBD | + | - | - | 210×210×240 | Heterogeneous N-glycan 1 | 1000 |
| HT2_WoBAT_WRBD | + | - | + | 210×210×255 | Heterogeneous N-glycan 2 | 1000 |
| HT2_WoBAT_WoRBD | + | - | - | 210×210×240 | Heterogeneous N-glycan 2 | 1000 |
| HTHS_WoBAT_WRBD | + | - | + | 210×210×255 | Heterogeneous N-glycan high sialic acid | 1000 |
| HTHS_WoBAT_WoRBD | + | - | - | 210×210×240 | Heterogeneous N-glycan high sialic acid | 1000 |
| HMC_RBD | - | - | + | 125×125×125 | Homogeneous complex N-glycan | 1000 |

**Table S2.** Interaction between the N322 glycan with the putative binding site in the RBD.

| System name | Hydrogen bonds |  | Weak hydrogen bonds |  | Hydrophobic interactions |  |
| --- | --- | --- | --- | --- | --- | --- |
|  | Mono A | Mono B | Mono A | Mono B | Mono A | Mono B |
| HMC_WBAT_WRBD | 0.2(0.8<sup>a</sup>) | 4.6(3.1) | 0.4(1.5) | 8.2(5.4) | 0.2(1.0) | 3.1(2.6) |
| HMC_WoBAT_WRBD | 3.2(2.1) | 5.3(3.0) | 8.3(4.0) | 8.8(4.8) | 4.3(2.7) | 3.8(2.8) |
| HMHM_WoBAT_WRBD | 0.9(1.8) | 4.1(2.1) | 1.1(2.1) | 6.8(3.6) | 0.4(0.9) | 3.7(2.9) |
| HT_WoBAT_WRBD | 4.2(2.9) | 3.1(2.4) | 7.4(4.6) | 6.2(3.7) | 3.9(2.9) | 4.4(3.2) |
| HT2_WoBAT_WRBD | 1.1(1.4) | 2.9(2.2) | 2.2(2.5) | 6.1(3.8) | 1.9(2.6) | 4.1(3.1) |
| HTHS_WoBAT_WRBD | 4.1(3.1) | 0.4(1.2) | 7.0(4.6) | 0.8(1.8) | 3.8(2.9) | 0.6(1.5) |

<sup>a</sup> standard deviation.

**Table S3.** The lipid composition used in this study for the membrane bilayer.

|  | CHOL (%) | POPC (%) | POPE (%) | POPS (%) | PSM (%) |
| --- | --- | --- | --- | --- | --- |
| Upper leaflet | 33 | 33 | - | - | 34 |
| Lower leaflet | 20 | 35 | 25 | 20 | - |

